# Amyloid pathology disrupts gliotransmitter release in astrocytes

**DOI:** 10.1101/679860

**Authors:** Anup G. Pillai, Suhita Nadkarni

## Abstract

Accumulation of amyloid-β peptide (Aβ), a hallmark of Alzheimer’s disease (AD), is associated with synchronous hyperactivity and dysregulated Ca^2+^ signaling in hippocampal astrocytes. However, the consequences of altered Ca^2+^ signaling on the temporal dynamics of Ca^2+^ and gliotransmitter release events at astrocytic microdomains are not known. We have developed a detailed biophysical model of microdomain signaling at a single astrocytic process that accurately describes key temporal features of Ca^2+^ events and Ca^2+^-mediated kiss-and-run and full fusion exocytosis. Using this model, we ask how aberrant plasma-membrane Ca^2+^ pumps and mGluR activity, molecular hallmarks of Aβ toxicity that are also critically involved in Ca^2+^ signaling, modify astrocytic feedback at a tripartite synapse. We show that AD related molecular pathologies increase the rate and synchrony of Ca^2+^ and exocytotic events triggered by neuronal activity. Moreover, temporal precision between Ca^2+^ and release events, a mechanism indispensable for rapid modulation of synaptic transmission by astrocytes, is lost in AD astrocytic processes. Our results provide important evidence on the link between AD-related molecular pathology, dysregulated calcium signaling and gliotransmitter release at an astrocytic process.

## Introduction

Bidirectional communication between neurons and astrocytes is crucial for normal brain function^1–5^. Despite the controversial role of astrocytes in information processing, it is well established that neuronal activity drives astrocytic Ca^2+^ signaling and concomitant release of gliotransmitters^6–8^. In parallel, disrupted astrocytic signaling has been implicated in several pathological conditions including cognitive decline and memory loss in AD^9,10^. Both *in vivo* and *in vitro* studies further confirm that accumulation of amyloid-β (Aβ), a pathological hallmark of AD, disrupts astrocytic Ca^2+^ signaling and astrocytic glutamate release^10–15^. Indeed, elevated resting Ca^2+^ levels and enhanced spontaneous and synchronous Ca^2+^ events have been observed *in vivo* in AD astrocytes^14^. Despite this evidence, the causal relationship between amyloid pathology, astrocytic Ca^2+^ hyperactivity and its downstream effects on gliotransmitter release at a single process has not been examined.

Ca^2+^ excitability in astrocytes primarily depends on metabotropic glutamate receptor (mGluR)-mediated generation of inositol-3-phosphate (IP_3_) and subsequent activation of IP_3_ receptors that release Ca^2+^ from the endoplasmic reticulum (ER)^7^. Enhanced mGluR5 expression and high ER to cytosol volume ratio of peripheral astrocyte processes (PAPs) that surround majority of cortical synapses allow fast and high-amplitude Ca^2+^ transients in these somewhat isolated glial compartments^16–25^. This specialized design together with presence of multiple Ca^2+^ sensors, vesicle types and recycling rates can trigger fast synchronous and slow asynchronous gliotransmitter release from individual astrocytic processes^26,27^. Measurements of individual exocytotic events indicate that mGluR stimulation leads to rapid kiss-and-run release of resident (docked) vesicles followed by slower full collapse fusions of newly recycled vesicles^22^. Consistent with this observation, several synaptotagmins (*Syts*) associated with exocytosis in neurons, including *Syt4 and Syt7,* have been reported in astrocytes^28,29^. *Syt4* structurally resemble *Syt1,* the fast neuronal Ca^2+^ sensor, and regulate rapid kiss-and-run fusions in astrocytes^29^, while *Syt7,* the neuronal asynchronous release sensor^30^, is largely associated with slow full fusion collapse of a vesicle^31–33^.Equipped with this elaborate machinery for exocytosis that is analogous to the presynaptic terminal in neurons, individual Ca^2+^ events in astrocytic microdomains exhibit precise spatiotemporal correlation with kiss-and-run release of a resident (docked) vesicle^22^. This correspondence between gliotransmitter release and calcium events is an integral part of synaptic transmission at a ‘tripartite synapse’ and critical for synchronization of neuronal activity^8,34^.

CA3 to CA1 synapses in the hippocampus are a potential neural substrate for various forms of learning and memory storage^35^. Encoding information in these synapses is input and synapse specific^36,37^. Anatomical and imaging studies show that astrocytic processes are in close proximity with most synapses in the hippocampus^19,38,39^. These findings together with their ability to generate calcium-induced gliotransmitter release suggest astrocytic processes as the critical center that facilitates information processing at individual synapses^40^.

While extensive experimental data and computational studies exists on somatic Ca^2+^ dynamics in astrocytes, a quantitative formulation of Ca^2+^ dynamics at a single astrocytic process and its relationship to Ca^2+^ mediated kiss-and-run and full fusion exocytosis has not been described^41–44^. We have developed a detailed biophysical model that incorporates multiple Ca^2+^ and IP_3_ signaling pathways alongside an elaborate kinetic scheme for synaptotagmins and vesicle recycling. The model reproduced Ca^2+^ and gliotransmitter release events at a single astrocytic process surrounding a typical CA3-CA1 synapse and is in quantitative agreement with experimentally observed Ca^2+^ and vesicular release events evoked by mGluR stimulation^22^. Using this modeling framework, we predict calcium activity and gliotransmitter releases at a single PAP in response to a range of presynaptic neurotransmitter release rates as well as characterize the consequences of AD-related molecular alterations in astrocytic signaling. Our results provide novel insights towards understanding how dysregulated astrocytic Ca^2+^ and gliotransmitter release can lead to aberrant synaptic signaling at a tripartite synapse in AD^45^.

## Methods

The calcium dynamics described by equations 1 and 2 combine fluxes (*Jx*) from various Ca^2+^ sources and Ca^2+^ regulating mechanisms that are present in the cytosol and ER, respectively (Fig. 1a). Equation 3 describes the production and degradation pathways of IP_3_, the intracellular signaling messenger. In the next sections we provide a detailed description of all Ca^2+^ and IP_3_ fluxes. Model parameters for all the components are in Tables 3-5.

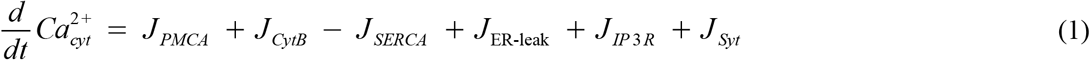

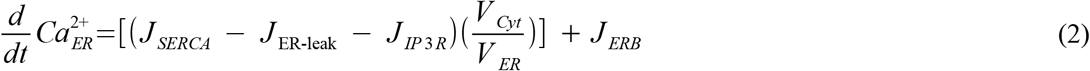

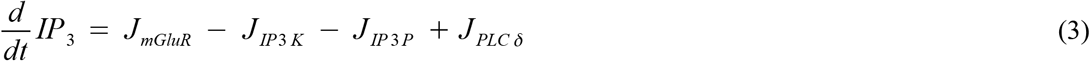

**Figure 1.**
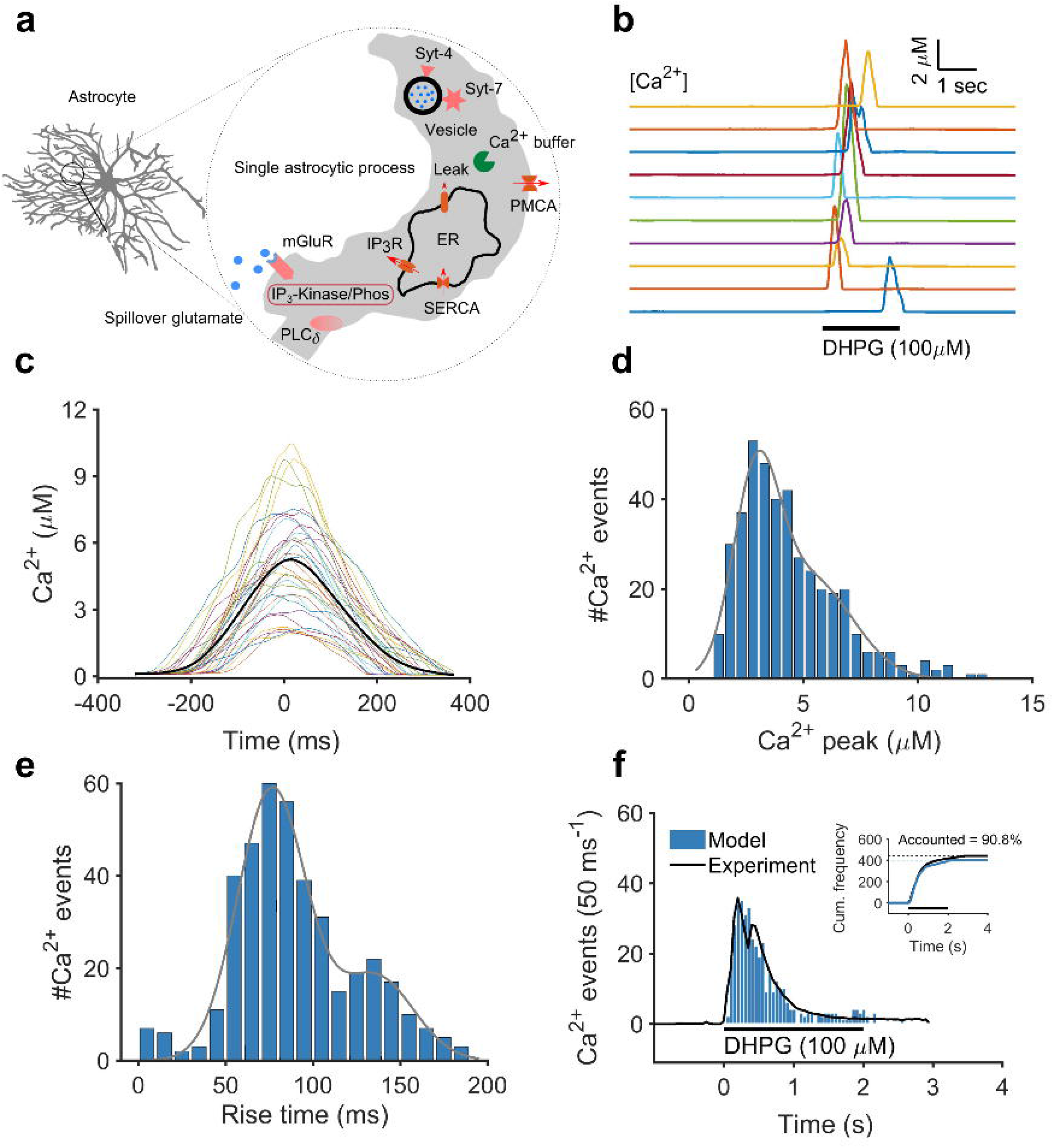
Description of the Ca^2+^ model and characterization of mGluR-mediated Ca^2+^ events. **a** Schematic of the single astrocyte process model with all the molecular components for Ca^2+^ signaling. **b** Representative Ca^2+^ responses from a set of independent trials of an astrocytic process stimulated with DHPG(100 μM, 2 sec). **c** Overlay of individual Ca^2+^ events aligned to the peak. **d** Peak amplitude histogram of Ca^2+^ events from a set of 400 independent trials that reproduced experimental data (f). **e** Distribution of Ca^2+^ event rise times. **f** Comparison of mGluR-mediated Ca^2+^ event histograms from the model and experimental data (Marchaland et. al^22^). *Inset* Cumulative distribution of Ca^2+^ events from the model and experiment, indicating that the model captures more than 90% of the all events. Gray lines indicate Gaussian fits.

### Calcium dynamics

The model includes multiple Ca^2+^ regulating components that are present on the plasma membrane, cytoplasm and ER. Plasma membrane high affinity Ca^2+^ ATPases (PMCAs) have been implicated in the maintenance of both resting and activity-dependent Ca^2+^ levels in astrocytes^46^. A three-state kinetic model captured the fast action of 2000 PMCAs, whereas slow action of cytosolic and ER Ca^2+^ buffers were modelled with two-state reaction equations (Tables 1 & 2). The parameters of cytosolic Ca^2+^ buffers (conc: 50 μM; K_D_: 20 μM) were adjusted to reproduce the fast kinetics of mGluR-dependent Ca^2+^ transients in processes as reported by previous studies^47^. We assumed low endogenous Ca^2+^ buffering in individual astrocytic processes similar to what has been observed in dendritic spines that promotes fast and high amplitude transients^48^. Apart from PMCAs and Ca^2+^ buffers, sarcoplasmic/ER Ca^2+^ ATPases (SERCAs) also have a key role in maintaining resting Ca^2+^ levels in both cytosol and ER. A Hill equation (Table 1) captured the slow SERCA activity whose Ca^2+^ pumping rate was adjusted to match the ER lumen Ca^2+^ concentration (100-800 μM)^49^. As a result, ER resting Ca^2+^ level (~ 400 μM) and pumping rate (250 μM s^−1^) in the model are higher compared to values in previous theoretical studies, 2-10 μM and ~ 1 μM s^-1^, respectively^41,50,51^. These considerations were important to reproduce the fast and high amplitude Ca^2+^ transients found at processes that primarily arise from stochastic opening of IP_3_-receptors on the ER membrane^22,23^. The ER membrane has a single cluster of 5 IP_3_R channels to resemble the patchy staining of IP_3_R_2_ on ER cisternae that generates the rapid Ca^2+^ puffs as reported by previous studies^16,52^. The model reliably captured the stochastic Ca^2+^ transients using a Langevin approximation of the Li-Rinzel IP_3_R model (Table 1) developed by Shuai et. al^51^. A passive Ca^2+^ leak channel maintained steady state ER Ca^2+^ level, although its precise role, despite its association with AD-related mutations, is not fully understood^53^.

**Table 1.** Ca^2+^ fluxes in the model

| Component name | Equation |
| --- | --- |
| PMCA<sup>68</sup> | $J_{PMCA} = (-M_0 Ca_{cyt}^{2+} k_{f1}) + (M_1 k_{b1}) + (M_0 k_l)$ $\frac{d}{dt} M_0 = (-M_0 Ca_{cyt}^{2+} k_{f1}) + (M_1 k_{b1}) + (M_2 k_{f3})$ $\frac{d}{dt} M_1 = (-M_1 k_{b1}) + (M_0 Ca_{cyt}^{2+} k_{f1}) + (-M_1 k_{f2})$ $\frac{d}{dt} M_2 = (-M_2 k_{f3}) + (M_1 k_{f2})$ |
| Cytosol Ca<sup>2+</sup> buffer<sup>100</sup> | $J_{CytB} = -Ca_{cyt}^{2+} C_0 k_f + C_1 k_b$ $\frac{d}{dt} C_0 = (-C_0 Ca_{cyt}^{2+} k_f) + (C_1 k_b)$ $\frac{d}{dt} C_1 = (-C_1 k_b) + (C_0 Ca_{cyt}^{2+} k_f)$ |
| SERCA<sup>101</sup> | $J_{SERCA} = V_{max} \frac{(Ca_{cyt}^{2+})^2}{(Ca_{cyt}^{2+})^2 + (k_a)^2}$ |
| ER-leak<sup>50</sup> | $J_{ER-leak} = k_{leak} (Ca_{ER}^{2+} - Ca_{cyt}^{2+})$ |
| IP<sub>3</sub> receptor<sup>51</sup> | $J_{IP3R} = v_{max} m_{\infty}^3 n_{\infty}^3 h^3 (Ca_{ER}^{2+} - Ca_{cyt}^{2+})$ $\frac{d}{dt} h = \alpha_h (1-h) - \beta_h h + \zeta(t)$ $\langle \zeta(t) \rangle = 0$ $\langle \zeta(t) \zeta(t') \rangle = \frac{(\alpha_h (1-h) + \beta_h h)}{n} \delta(t-t')$ $d_i = \frac{b_i}{a_i}, \quad q_2 = d_2 \left( \frac{IP_3 + d_1}{IP_3 + d_3} \right)$ $n_{\infty} = \frac{IP_3}{IP_3 + d_1}, \quad m_{\infty} = \frac{Ca_{cyt}^{2+}}{Ca_{cyt}^{2+} + d_5}$ $\alpha_h = a_2 d_2 \frac{(IP_3 + d_1)}{(IP_3 + d_3)}, \quad \beta_h = a_2 Ca_{cyt}^{2+}$ |
| ER Ca<sup>2+</sup> buffer<sup>102</sup> | $J_{ERB} = -Ca_{ER}^{2+} B_0 k_f + B_1 k_b$ $\frac{d}{dt} B_0 = (-B_0 Ca_{ER}^{2+} k_f) + (B_1 k_b)$ $\frac{d}{dt} B_1 = (-B_1 k_b) + (B_0 Ca_{ER}^{2+} k_f)$ |
| Synaptotagmin 4 & 7<sup>68,69</sup> | $J_{Syt} = J_{Syt4} + J_{Syt7}$ $J_{Syt4} = (-2 S_0 k_{f4} Ca_{cyt}^{2+}) + (S_1 k_{b4}) + (-S_1 k_{f4} Ca_{cyt}^{2+}) + (2 S_2 k_{b4} b_4)$ $J_{Syt7} = + (-5 Y_0 k_{f7} Ca_{cyt}^{2+}) + (Y_1 k_{b7}) + (-4 Y_1 k_{f7} Ca_{cyt}^{2+}) + (2 Y_2 k_{b7} b_7)$ $+ (-3 Y_2 k_{f7} Ca_{cyt}^{2+}) + (3 Y_3 k_{b7} b_7^2) + (-2 Y_3 k_{f7} Ca_{cyt}^{2+}) + (4 Y_4 k_{b7} b_7^3)$ $+ (-Y_4 k_{f7} Ca_{cyt}^{2+}) + (5 Y_5 k_{b7} b_7^4)$ |

**Table 2.** IP_3_ fluxes in the model

| Component name | Equation |
| --- | --- |
| mGluR<sup>103</sup> | $J_{mGluR} = V_{max} \frac{(Glu)^2}{(Glu)^2 + (k_a)^2}$ |
| IP<sub>3</sub> 3-kinase<sup>54</sup> | $J_{IP_3 K} = V_{max} \frac{(Ca_{cyt}^{2+})^4}{(Ca_{cyt}^{2+})^4 + (k_{a1})^4} \times \frac{IP_3}{IP_3 + k_{a2}}$ |
| IP<sub>3</sub> 5-phosphatase<sup>44,103</sup> | $J_{IP_3 P} = V_{max} (IP_3 - IP_3^{base})$ |
| PLC<sub>δ</sub><sup>44</sup> | $J_{PLC\delta} = \frac{V_{max}}{(1 + (IP_3/k_{a1}))} \times \frac{Ca_{cyt}^{2+}}{(Ca_{cyt}^{2+} + k_{a2}^2)}$ |

**Table 3.** Model parameters for Ca^2+^ and IP_3_ dynamics

| Component name | Parameters |
| --- | --- |
| PMCA<sup>68</sup> | $k_{f1} = 15 \mu M^{-1} s^{-1}$, $k_{b1} = 20 s^{-1}$, $k_{f2} = 20 s^{-1}$, $k_{f3} = 100 s^{-1}$, $k_l = 0.6 s^{-1}$, *PMCA density* = $400 \mu m^{-2}$ |
| Cytosol Ca<sup>2+</sup> buffer<sup>100</sup> | $k_f = 60 \mu M^{-1} s^{-1}$, $k_b = 1200 s^{-1}$,<br>*buffer concentration* = $50 \mu M$ |
| SERCA<sup>101</sup> | $V_{max} = 250 \mu M s^{-1}$, $k_a = 100 nM$, $n = 2$ |
| ER-leak | $k_l = 0.2 s^{-1}$ |
| IP<sub>3</sub> receptor<sup>51</sup> | $a_1 = 400 \mu M s^{-1}$, $a_2 = 0.2 \mu M^{-1} s^{-1}$, $a_3 = 400 \mu M^{-1} s^{-1}$, $a_4 = 0.2 \mu M^{-1} s^{-1}$, $a_5 = 20 \mu M^{-1} s^{-1}$,<br>$b_1 = 52 s^{-1}$, $b_2 = 0.2 s^{-1}$, $b_3 = 377.36 s^{-1}$, $b_4 = 0.02 s^{-1}$, $b_5 = 1.64 s^{-1}$, $v_{max} = 6.0 s^{-1}$, $n = 5$ |
| ER Ca<sup>2+</sup> buffer<sup>102</sup> | $k_{f1} = 1 \mu M^{-1} s^{-1}$, $k_{b1} = 80 s^{-1}$, *buffer concentration* = $10 \mu M$ |
| mGluR<sup>103</sup> | $V_{max} = 0.65 \mu M s^{-1}$, $k_a = 6 \mu M$, $n = 2$ |
| IP<sub>3</sub> 3-kinase<sup>54</sup> | $V_{max} = 5 \mu M s^{-1}$, $k_{a1} = 0.4 \mu M$, $n_1 = 4$, $k_{a2} = 10 \mu M$, $n_2 = 1$ |
| IP<sub>3</sub> 5-phosphatase<sup>44,103</sup> | $V_{max} = 1.25 s^{-1}$ |
| PLC<sub>δ</sub><sup>44</sup> | $V_{max} = 0.02 \mu M s^{-1}$, $k_{a1} = 1 \mu M$, $n_1 = 2$, $k_{a2} = 1.5 \mu M$ |

**Table 4.**
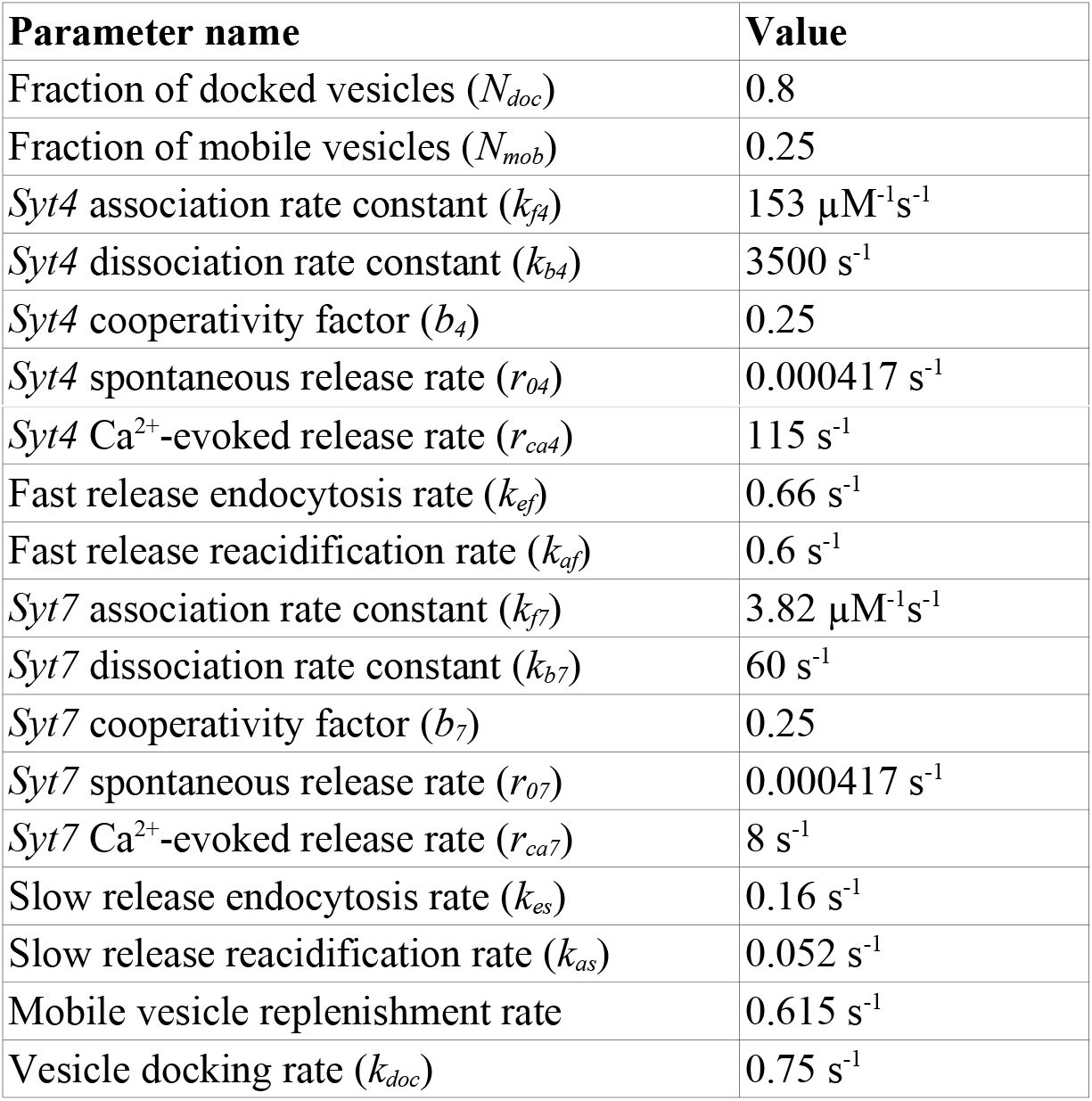
Model parameters for vesicular release (Adapted from multiple studies^22,66,68,69^)

**Table 5.**
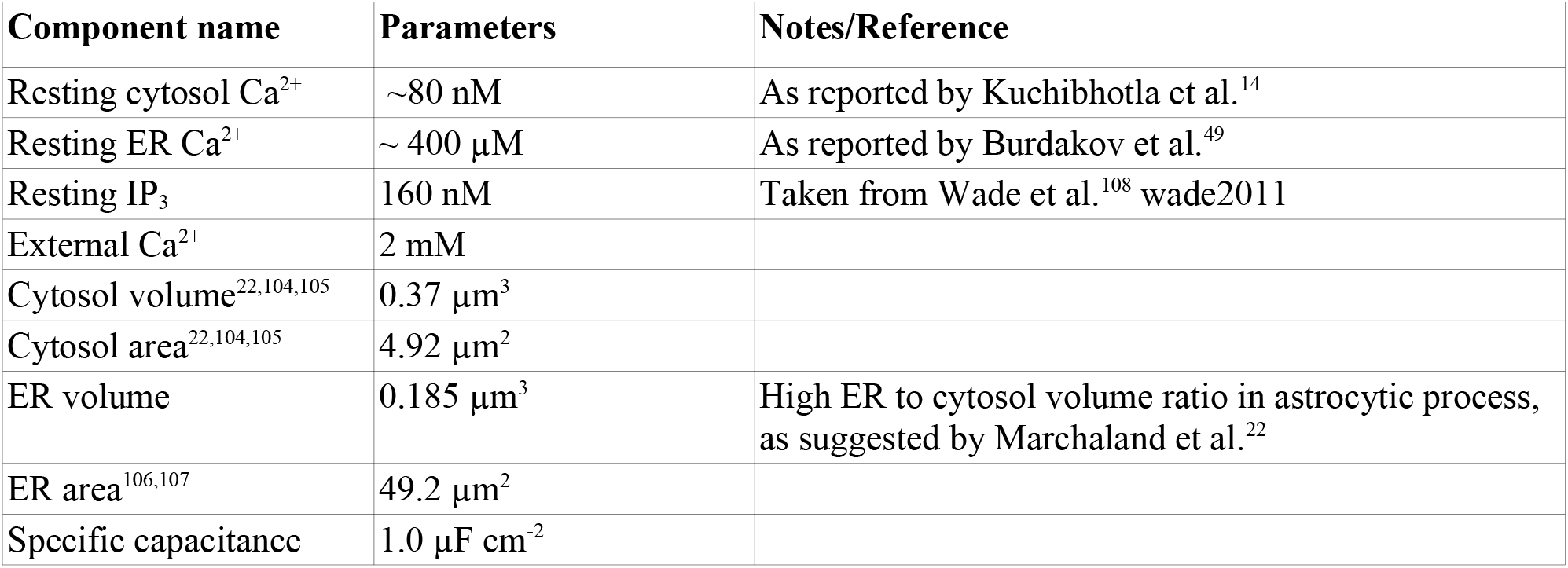
Additional parameters/details of the model

| Component name | Parameters | Notes/Reference |
| --- | --- | --- |
| Resting cytosol Ca<sup>2+</sup> | ~80 nM | As reported by Kuchibhotla et al.<sup>14</sup> |
| Resting ER Ca<sup>2+</sup> | ~ 400 $\mu\text{M}$ | As reported by Burdakov et al.<sup>49</sup> |
| Resting IP<sub>3</sub> | 160 nM | Taken from Wade et al.<sup>108</sup> wade2011 |
| External Ca<sup>2+</sup> | 2 mM |  |
| Cytosol volume<sup>22,104,105</sup> | 0.37 $\mu\text{m}^3$ | |
| Cytosol area<sup>22,104,105</sup> | 4.92 $\mu\text{m}^2$ | |
| ER volume | 0.185 $\mu\text{m}^3$ | High ER to cytosol volume ratio in astrocytic process, as suggested by Marchaland et al.<sup>22</sup> |
| ER area<sup>106,107</sup> | 49.2 $\mu\text{m}^2$ | |
| Specific capacitance | 1.0 $\mu\text{F cm}^{-2}$ | |

### IP_3_ dynamics

While the major IP_3_ generation pathway is through the activation of mGluRs by glutamate spillover from adjacent presynaptic terminals, IP_3_ is also generated in a Ca^2+^-dependent manner through PLC_δ_ activity. IP_3_ degradation in astrocytes are thought to be mediated by two separate molecular pathways, namely, inositol polyphosphate 5-phosphatase and IP_3_ 3-kinase, the former operates via a Ca^2+^-independent mechanism while the latter is Ca^2+^ activated^54^. The model approximated all these slow IP_3_ regulation mechanisms using four separate Hill equations (Table 2).

### Gliotransmitter release

We modelled Ca^2+^-dependent gliotransmitter release using detailed kinetics schemes for two distinct synaptotagmins (*Syt4* and *Syt7*) as well as the pathways of endocytosis, recycling and vesicle docking (Fig. 2a and supplementary Fig. 5). *Syt4,* has low Ca^2+^ affinity (K_D_: 22 μM) and fast forward reaction rates compared to *Syt7. Syt4* can therefore be activated in tight spaces close to the plasma membrane and ER where the vesicles are docked and Ca^2+^ transients are fast and have high amplitudes^22^. In comparison, *Syt7* has high Ca^2+^ binding affinity (~ 15 μM) but slow binding rates. Low amplitude and but long lasting Ca^2+^ rises can therefore activate *Syt7.* As a consequence, Ca^2+^ events that are generated far from a vesicle can still contribute to the overall gliotransmitter release.

**Figure 2.**
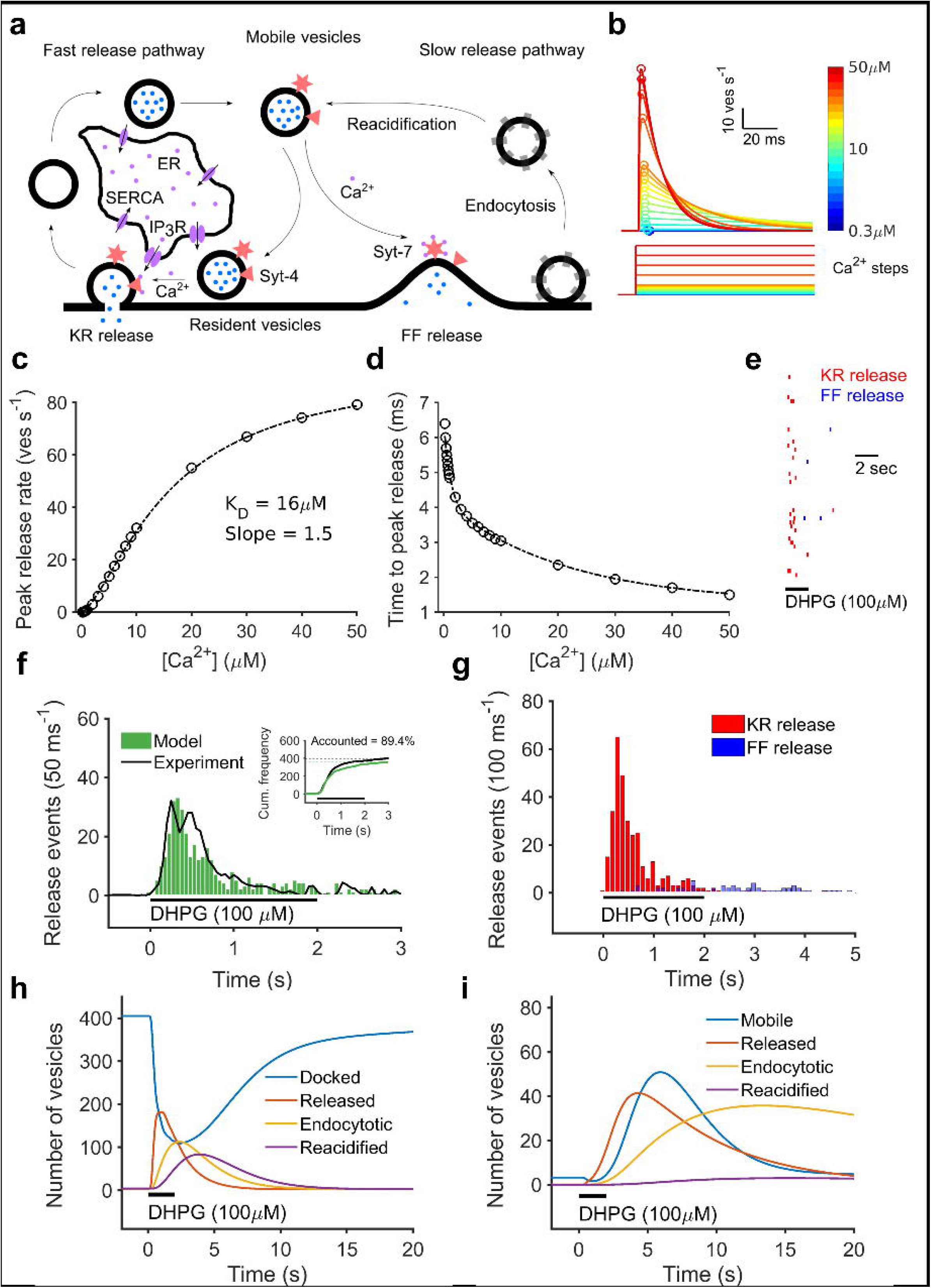
The gliotransmsitter release model and characterization of mGluR-mediated release events. **a** Schematic of the gliotransmitter release model, indicating the two synaptotagmins (*Syt4* & *Syt7*) that independently trigger kiss-and-run and full fusion releases from docked (resident) and mobile vesicles, respectively. **b** Total gliotransmitter release rate from an astrocytic process whose intracellular Ca^2+^ is clamped at different concentration steps. **c** Ca^2+^ dependence of peak release rate is fitted with a Hill equation. **d** Time of peak release rate decrease exponentially with increasing cytosolic Ca^2+^ levels. **e** Perievent raster plot of mGluR-mediated kiss-and-run and full fusion events from a subset of simulation trials. **f** Comparison of temporal histogram of mGluR-mediated release events collected from 400 independent simulation trials and experimental data (Marchaland et al.^22^). *Inset* Cumulative histogram of gliotransmitter release from the single process model versus experimental data (Marchaland et al.^22^). **g** Distribution of kiss-and-run and full fusion releases from 400 independent trials of a single astrocyte process stimulated with DHPG (100 μM, 2 sec). **h** Kinetics of release, endocytosis, and reacidification of docked vesicles after stimulation with DHPG. **i** Kinetics of release, endocytosis, and reacidification of mobile vesicles.

Consistent with the difference in Ca^2+^ binding kinetics of the two synaptotagmins, *Syt4* and *Syt7* coordinate distinct modes of release from different vesicle pools. *Syt4* is associated with rapid kiss-and-run exocytosis of the docked vesicles close to the juxtaposition between ER and plasma membrane, whereas *Syt7* is associated with slow full fusion releases^28,29,33,55^. Interestingly, mobile vesicles that are ‘not’ localized in close spaces where Ca^2+^ concentration is high have been observed in astrocytic processes^56^. These vesicles are likely to be newly recycled and not docked and their release can be attributed to full fusion via *Sty7.* In summary, high affinity binding sites in *Syt7* are activated by low amplitude Ca^2+^ elevations and promote slow full fusions of mobile vesicles, while docked vesicles are exocytosed in kiss-and-run mode by *Syt4.* Each of these pathways also follow distinct rates of endocytosis and recycling and have been modelled accordingly^22^.

We have systematically collated data from diverse experimental studies to refine t h e model parameters^22,24,57,58^. Despite differences in experimental settings, temporal release histograms collected from these multiple studies on gliotransmitter release exhibited striking similarities in rise time, peak and decay time (Supplementary Fig. 2). From the reported values of imaging area (~ 1115 μm^2^) and TIRF evanescent field (~ 100 nm) by these studies, we calculated an average imaging volume of roughly 111 μm^3^ per astrocyte. Together with previous measurements on total astrocytic volume (24,465 μm^3^) and the number of processes (100,000) in the CA1 layer of the rodent hippocampus, we estimated between 400-500 processes within the imaging volume^20,59^. Cumulative distributions of release histograms from these studies also indicated that a maximum of ~ 400 vesicles are released within this volume (Supplementary Fig. 3). We therefore, collected data from 400 independent simulation trials of a single astrocytic process to match the model results with experimental data. At resting state, 80% of the total vesicles are docked while the rest are in the mobile pool.

A typical chain of events starts with presynaptic release that increases extracellular glutamate concentration (~200 μM). We used a simple first order reaction equation (tau = 6.25 ms) to capture the rapid rise and decay (~ 10 ms) of glutamate levels near astrocytic processes^60^. Individual release events activated mGluRs to increase intracellular IP_3_ levels (~ 175 nM) that decayed to resting levels (~ 160 nM). The increase in IP_3_ caused cytoplasmic Ca^2+^ transients by stochastic opening of IP_3_Rs on the ER membrane (Supplementary Fig. 1). The change in Ca^2+^ level was small (5-20 nM) at low frequency of activation. However, at higher rates of presynaptic glutamate release, the rapid build of IP_3_ generated high amplitude Ca^2+^ events (5-10 μM) that activated synaptotagmins and caused concomitant gliotransmitter release events. We characterized the release machinery by clamping intracellular Ca^2+^ levels and measuring peak release (R) rate which was fitted to a Hill equation (4) with *Vmax, Kd* and *coeff* as free parameters.

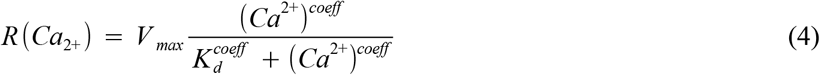

We additionally fitted events rates (*H*) for Ca^2+^ and gliotransmitter release with a Hill equation (Eq. 5) to quantify its relationship with stimulus train frequency (*f*). Where *Hmin, Hmax* and EC_50_ refer to minimum, maximum and half-max of event rate. All the parameters were fit recursively using the Nelder-Mead simplex algorithm.

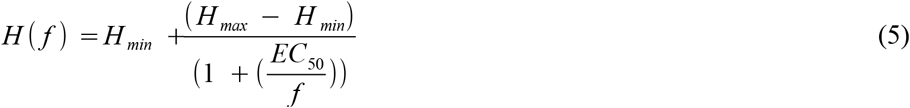

### Data analysis

#### Calcium responsiveness measure

We computed Ca^2+^ responsiveness for a single process by averaging Ca^2+^ signals across trials (N = 1000). Astrocytic process was stimulated with the spillover glutamate profile from neuronal release at different firing frequencies (0.1-100 Hz) that lasted for 30 secs. The responsive measure *r* was computed by normalizing the area under the curve as described in the below equation (Eq. 6), where *t_0_* and *T_stim_* refer to stimulation onset and duration, respectively.

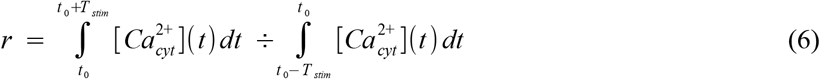

#### Synchrony measure

We computed synchrony of Ca^2+^ and release events from independent trials (N = 1000) of a single astrocytic process stimulated with the same protocol as explained above. Event times for Ca^2+^ transients corresponded to the peak time of all the Ca^2+^ spikes whose amplitudes were above 300 nM. Individual release events were obtained from Poisson distribution of the instantaneous release rate at each simulation time step (50 μs). For each stimulation frequency, we calculated the Pinsky-Rinzel measure of synchrony^61^ from the set of phases, *l_k_*(*j, m*) for every *m*^th^ event in *j*th trial (Eq. 7). Where k, 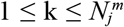, corresponds to all the events within the time interval 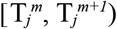 from all trials excluding the jth trial itself. The operation transforms event times to a corresponding value spanning from 0 to 1.

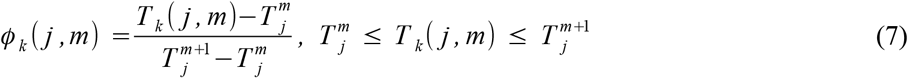

Synchrony was then calculated from the vector of phases using the Strogatz and Mirollo^62^ method as defined below (Eq. 8)

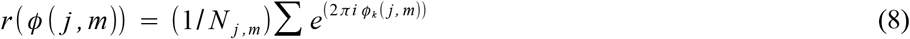

Overall synchrony for the set of N trials was obtained from the real part of r(l (j,m)) averaged across all the trials (*j*) and events (*m*). A value of 1 corresponds to a perfect alignment of event times across all the trials whereas 0 meant events arriving randomly.

#### Computation of correlation between Ca^2+^ and gliotransmitter release events

We quantified time-varying correlation between Ca^2+^ and gliotransmitter release events across independent trials using normalized joint peristimulus time histogram (nJPSTH), a method introduced by Aertsen at al. to compute correlations between neuronal spike trains^63^. For each trial of the stimulus protocol similar to the one described above for Ca^2+^ responsiveness, we computed peristimulus time histograms (PSTHs, 100 ms bins) for both Ca^2+^ and release events. For each non-zero bin of the PSTHs, the value of JPSTH matrix whose row and column indices correspond to the PSTH time bins of release and Ca^2+^ events, respectively, was incremented by one. The above procedure was repeated for all the 1000 trials. The JPSTH matrix was normalized by subtracting and dividing it with the matrix product and standard deviations, respectively, of normalized Ca^2+^ and release PSTHs. Cross-correlation was computed by summing the para-diagonal bins of the normalized JPSTH matrix from one second before to 11 seconds after the stimulus. Coincidence histogram was obtained from values along the main diagonal of the nJPSTH matrix. We quantified the relationship between peak cross-correlation across different stimulation frequencies by fitting to a logistic function as described below (Eq. 9). Where, *C_top_* and *C_slope_* refer to peak and steepness of the response, while *f_mid_* is the frequency at which response attains halfmaximum.

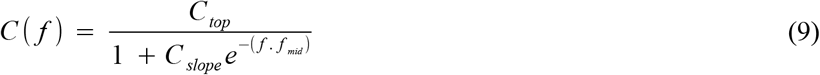

#### Numerical simulation

The simulations were run with 50 μs time step on the institution’s HPC facility (1500 nodes, PBSPro scheduler). The data was analyzed using custom Python scripts (www.python.org).

## Results

### Model of Ca^2+^ signaling at a single astrocytic process

A schematic of the Ca^2+^ model is illustrated in figure 1a. The model has two coupled compartments for Ca^2+^ dynamics, (1) an intracellular cytoplasmic space and (2) ER compartment. PMCA pumps on the plasma membrane mediate slow bidirectional Ca^2+^ flow, whereas rapid Ca^2+^ elevations within a process primarily arise from stochastic opening of IP_3_ receptors present on the ER membrane. The kinetics of Ca^2+^ spikes are further shaped by Ca^2+^ buffers and SERCAs. The model captures stochastic Ca^2+^ dynamics in single astrocytic processes upon mGluR stimulation (details of each component are in the Methods section).

The study by Marchaland et al. reported on the characteristics of Ca^2+^ events from single astrocytic processes activated by DHPG (100 μM, 2 secs), a highly specific group 1 mGluR agonist^22^. Using the same experimental paradigm, we first examined Ca^2+^ event characteristics from 400 independent runs of the single process model. Representative time courses and overlay of stochastic Ca^2+^ spikes from a subset of trials are presented in figures 1b and c. Peak amplitude, rise time and decay constant of the events were normally distributed with peaks at 2.4 μM, 80 ms and 120 ms, respectively (Figs. 1d & e). In good agreement with the experimental results, we find that the temporal distribution of Ca^2+^ events decayed exponentially (τ = 378 ms) (Fig. 1f). The model’s close match with experimental result is further evident from the close overlap (~ 90%) between Ca^2+^ event cumulative histograms (Fig. 1f, *inset*).

### Modeling vesicular release at a single astrocytic process

Despite several reports on Ca^2+^-dependent gliotransmitter release from astrocytes, a biophysical framework that links calcium rise at a single process to kiss-and-run and full fusion exocytosis is not present. Based on a broad understanding of astrocytic Ca^2+^-sensors (synaptotagmins), vesicle distribution and recycling times, we propose a detailed kinetic scheme for the astrocytic release machinery at a single process (Fig. 2a & Supplementary Fig. 5).

Two synaptotagmins, *Syt4 and Syt7,* have been reported in astrocytes^28,29,33,64^. *Syt4* has a single low affinity domain (C_2_B) that binds 2 Ca^2+^ ions with fast forward rates^31^. In contrast, *Syt7* has two calcium domains (C_2_A and C_2_B) that have high affinity but slow forward reaction rates^65^. This makes *Syt7* a slow calcium sensor that operates under low Ca^2+^ levels. TIRF imaging studies using fluorescently labelled vesicles also indicate fast kiss-and-run-like confined releases alongside slow spreading full fusions events^27,32^. It is also established that *Syt4* with a single Ca^2+^ binding domain can only promote kiss-and-run exocytosis^31^. Conversely, capacitance measurements in astrocytes that primarily reflects full fusion exocytosis estimated a Hill’s coefficient of 5 that match with the Ca^2+^ binding sites in *Syt7^66^*.

Based on these findings, we modeled two separate Ca^2+^ sensors, *Syt4* and *Syt7,* that have different Ca^2+^ affinities and independently mediate fast (kiss-and-run release) and slow (full fusion) gliotransmitter release respectively, in astrocytic process. *Syt4,* having low Ca^2+^ affinity and fast reaction kinetics, is primarily activated by sharp and high amplitude Ca^2+^ events that have been observed near ER tubuli. On the other hand, *Syt7,* with high Ca^2+^ affinity and slow kinetics, specifically respond to slow and low amplitude Ca^2+^ rises. Analogous to vesicle recycling at the presynaptic terminal^67^, we further assumed that every vesicle once released is subsequently endocytosed and transported (*mobile*) to a docking site (Fig. 2a). Separately, it has also been suggested that kiss-and-run and full fusion may arise from discrete vesicle pools^22,27^. Estimates on release parameters including release rate, vesicles per process and rates of endocytosis and vesicle recycling were obtained from published literature and are detailed in Tables 1-5.

Similar to what has been studied at the presynaptic terminal, we first examined the relationship between Ca^2+^ and peak release rate at a single process^68,69^. Clamping intracellular Ca^2+^ transiently increased release rate that decayed back to zero in tens of milliseconds (Fig. 2b). Estimates on the release machinery parameters, *K_D_* and binding sites (*slope),* were obtained by fitting the peak release rate to a Hill equation (Eq. 4; Fig. 2c). Decay of time-to-peak release was fit with a bi-exponential function with two time constants (Fig. 2d).

We next examined gliotransmitter release from 400 independent processes to validate the model results with previous experimental studies that measured single exocytotic events from entire astrocytes^22,24,57,58^. Similar to the study by Marchaland et al., we stimulated astrocytic process with DHPG (100 μM, 2 secs) to evoke Ca^2+^ transients and concomitant gliotransmitter events^22^. A representative raster plot of kiss-and-run and full fusion events from a subset of astrocytic process stimulated with DHPG is shown in figure 2e. The model indicates rapid rise in release events that peaked at roughly 250 ms after the stimulus onset and decayed exponentially (Fig. 2f). Both temporal and cumulative release histograms (Fig. 2f *inset*) from the model are in good agreement with the study by Marchaland et al. The model further indicates that a majority of the vesicles are exocytosed immediately after the stimulus via kiss-and-run mode followed by slow and asynchronous full fusions (Fig. 2g). Total number of docked vesicles from all the processes decreased rapidly after DHPG application while the number of vesicles in the mobile pool increased. Similar to what has been observed experimentally, the release, endocytosis and reacidification of docked vesicles are faster compared to mobile vesicles (Figs. 2h-i)

### Modeling astrocytic processes with AD pathology

mGluR signaling and PMCA pump activity, two critical regulators of astrocytic calcium signaling, are modified in AD. Specifically, mGluRs responses are enhanced in AD astrocytes^13,70,71^, while Aβ inhibits Ca^2+^ signaling through PMCAs^72–74^.

Similar what has been observed experimentally, we up-regulated mGluR signaling in the AD-mGluR group by increasing its affinity for glutamate (K_D_, control: 6 μM, AD-mGluR: 3 μM) and peak IP_3_ production rate (*V_max_,* control: 0.65 μMs^−1^, AD-mGluR: 1.4 μMs^−1^). As expected, peak IP_3_ response to constant glutamate stimulation lasting 30 sec is both higher and left shifted in the AD-mGluR group in comparison to the control group, indicating an overall increase in IP_3_ production (Fig. 3a).

**Figure 3.**
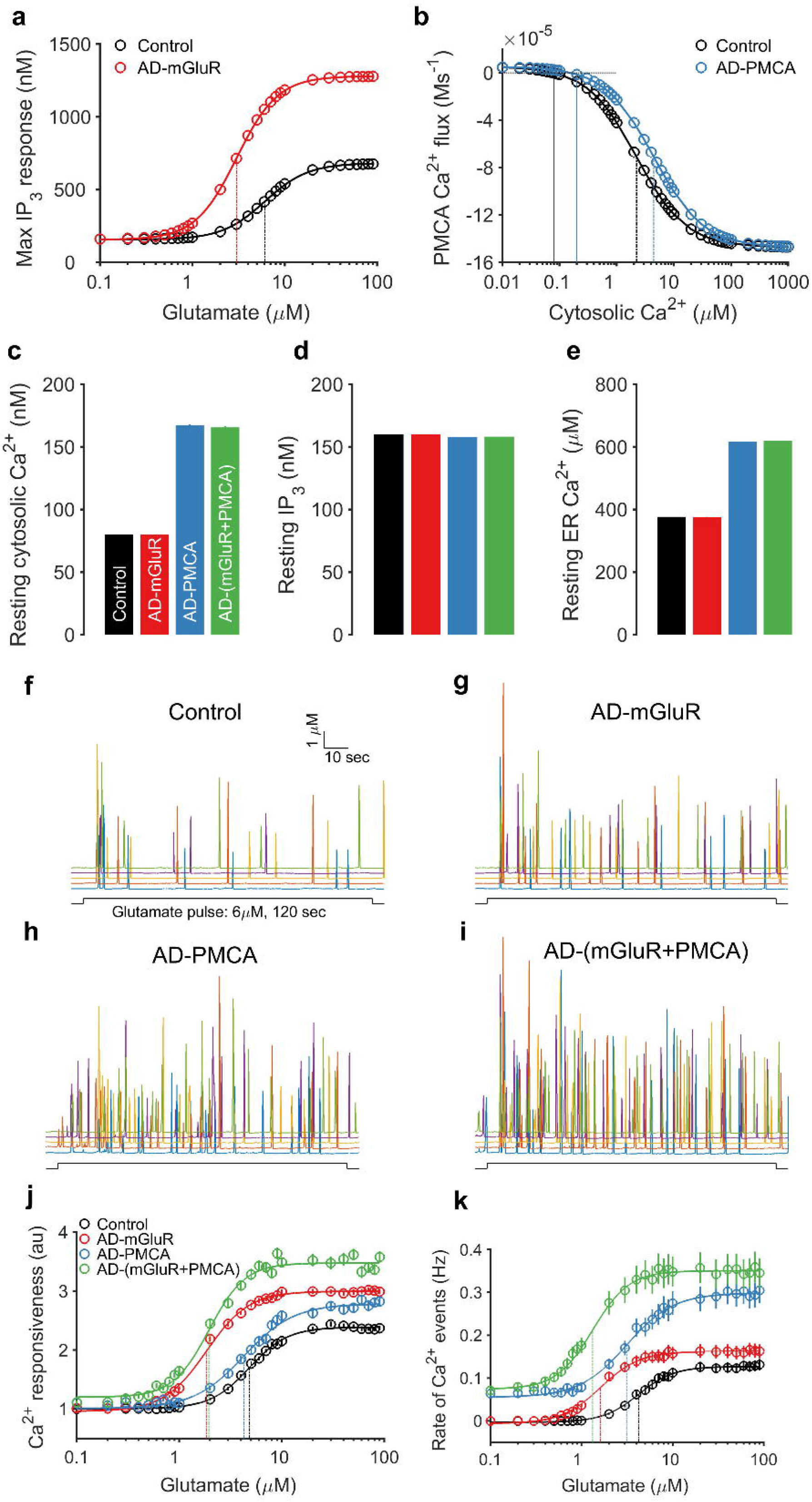
A D related alterations in mGluR and PMCA signaling modifies Ca^2+^ activity of astrocytic process at resting state and when stimulated by glutamate. **a** Dose response curve for mGluR-dependent IP_3_ production is shifted leftward in the AD-mGluR astrocytic process. **b** Intracellular Ca^2+^ dependence of PMCA steady state Ca^2+^ flux is shifted to the right in the AD-PMCA astrocytic process. **c** Resting cytosolic Ca^2+^ level is higher in astrocytic process with impaired PMCA activity. **d** Resting IP_3_ levels are not affected by AD-related molecular signaling. **e** Resting ER Ca^2+^ level is enhanced in AD astrocytic processes with altered PMCA signaling. **f-i** Representative Ca^2+^ responses from control and AD astrocytic processes that are stimulated with a prolonged glutamate pulse (6 μM, 120 s). **j** Glutamate-induced Ca^2+^ responsiveness is enhanced in all AD astrocytic processes. **k** Ca^2+^ event rate during glutamate stimulation is higher in all AD astrocyte groups when compared to the control group. Solid lines are Hill equation fits and vertical dotted lines indicate glutamate concentrations at half-maximal response.

We next modeled the shift PMCA’s Ca^2+^ dependency in the AD-PMCA group by decreasing its affinity for Ca^2+^ (K_D_, control: 1.3 μM, AD-PMCA: 4 μM). In the AD-PMCA group, the decrease in PMCA Ca^2+^ affinity could be observed as a rightward shift in its steady-state outward Ca^2+^ flux when the level of intracellular Ca^2+^ was clamped to different concentrations (Fig. 3b). An interesting side effect of decreased PMCA efficiency is the rise in resting Ca^2+^ levels in both the ER and cytoplasm in the AD-PMCA astrocytic process (control: 80 nM, AD-PMCA: 166 nM) (Figs. 3c & e). However, resting IP_3_ levels where not affected in any of the AD groups (Fig. 3d).

We further probed the effect of PMCA and mGluR manipulation on astrocytic Ca^2+^ dynamics by stimulating control and all AD groups with a standard stimulus protocol used in several experimental studies that examined mGluR-mediated Ca^2+^ signaling in astrocytes^75,76^. Consistent with experimental results, prolonged glutamate application (6 μM, 120 s) induced Ca^2+^ spikes in a majority AD astrocytic process (Figs. 3f-i, colors represent independent trials). Ca^2+^ oscillations were quantified using the responsiveness measure which is a readout for cumulative response for the entire stimulus duration (See Model and Methods for details). All the AD astrocytic groups displayed higher responsiveness and Ca^2+^ event rate compared to control, this is evident from the leftward-shift in the dose-response curves (Figs. 3j-k). Interestingly, AD groups with impaired PMCA signaling exhibited spontaneous Ca^2+^ events even at a very low glutamate concentration (< 1 μM) where the control astrocytic process is silent (Fig. 3k).

### Ca^2+^ events are enhanced in astrocytic process with AD pathology

Several studies have reported increased Ca^2+^ activity in astrocytes from AD animal models and when treated with amyloid-β^11,77^. Additionally, astrocytes with AD pathology also exhibit unusually high spontaneous Ca^2+^ activity^14,77,78^. Exploring further on our previous results on astrocytic responses to constant glutamate pulses, we next quantified Ca^2+^ signaling in a n astrocytic process using a physiologically realistic stimulus with the same concentration and temporal profile of spillover glutamate from presynaptic release^60^ (details in materials and methods).

As representative examples we show Ca^2+^ transients evoked by spillover glutamate released at 3 Hz (Fig. 4a-d). Ca^2+^ events were more frequent and larger in amplitude in astrocytic process from all the AD groups compared to the control group. Additionally, in AD groups with impaired PMCAs, spontaneous events could be observed before the stimulus onset (*Dotted box* in Figs. 4c & d).

**Figure 4.**
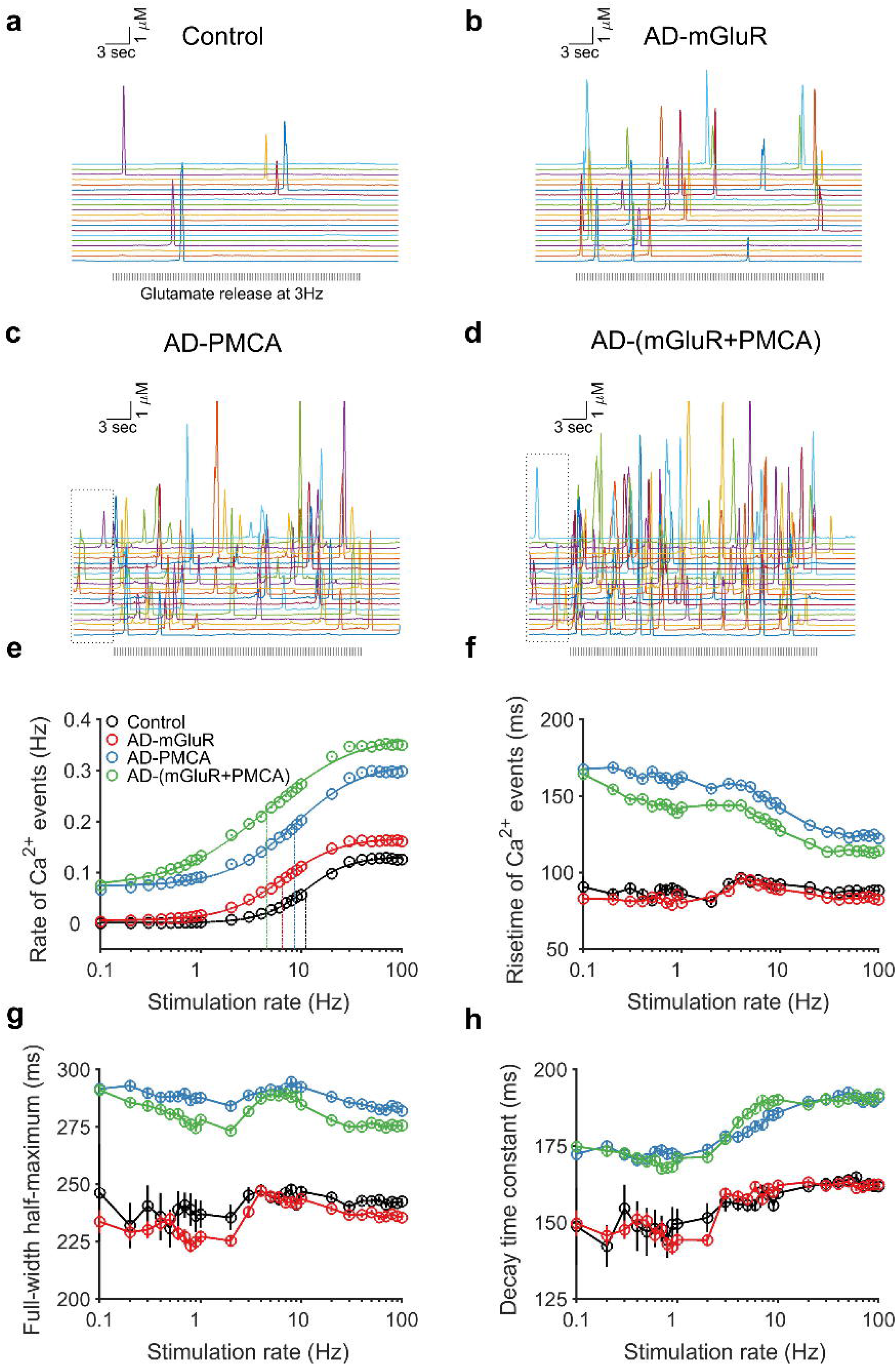
Spontaneous and evoked Ca^2+^ activity are increased in A D astrocytic processes with AD pathology. **a-d** Ca^2+^ transients recorded from control and AD astrocytic processes stimulated with spillover glutamate from presynaptic release at 3 Hz. Dotted boxes indicate the part where astrocytic process display spontaneous Ca^2+^ transients. **e** The rate of Ca^2+^ event rate increased non-linearly with stimulation frequency. Solid lines are fits to a Hill equation and dotted vertical lines indicate stimulation frequency for half-maximal response. Risetime (**f**), full-width at half-maximum (**g**) and decay time (**h**) of Ca^2+^ events averaged for all the events across 1000 independent simulation trials for different frequencies over the entire stimulation regime are lower in AD groups with aberrant PMCA, but not mGluR, signaling.

To further characterize these Ca^2+^ responses, we stimulated astrocytic processes at different presynaptic release rates (0.1-100 Hz) resembling the broad range of neuronal activity observed *in vivo* (Figs. 4e-h). Half maximums of Ca^2+^ event rate (EC_50_) obtained from fits to a Hill equation (Eq. 5) were low in all the AD groups compared to control astrocytic process (Fig. 4e), implying enhanced Ca^2+^ activity. Impairment in PMCA activity, apart from increasing spontaneous activity, also lowered rise time, full-width at half-maximum and decay time in all the stimulation regimes (Figs. 4f-h).

### Gliotransmitter release events are enhanced in astrocytic process with AD pathology

We next examined the impact of AD-induced alterations on Ca^2+^-mediated exocytosis. A single astrocytic process was stimulated (~ 1000 trials) with the temporal profile of spillover glutamate released at different rates (0.1-100 Hz). Representative raster plots (30 trials; 30 sec) of kiss-and-run and full fusion release from 3 Hz stimulation are described in figure 5 a-d.

**Figure 5.**
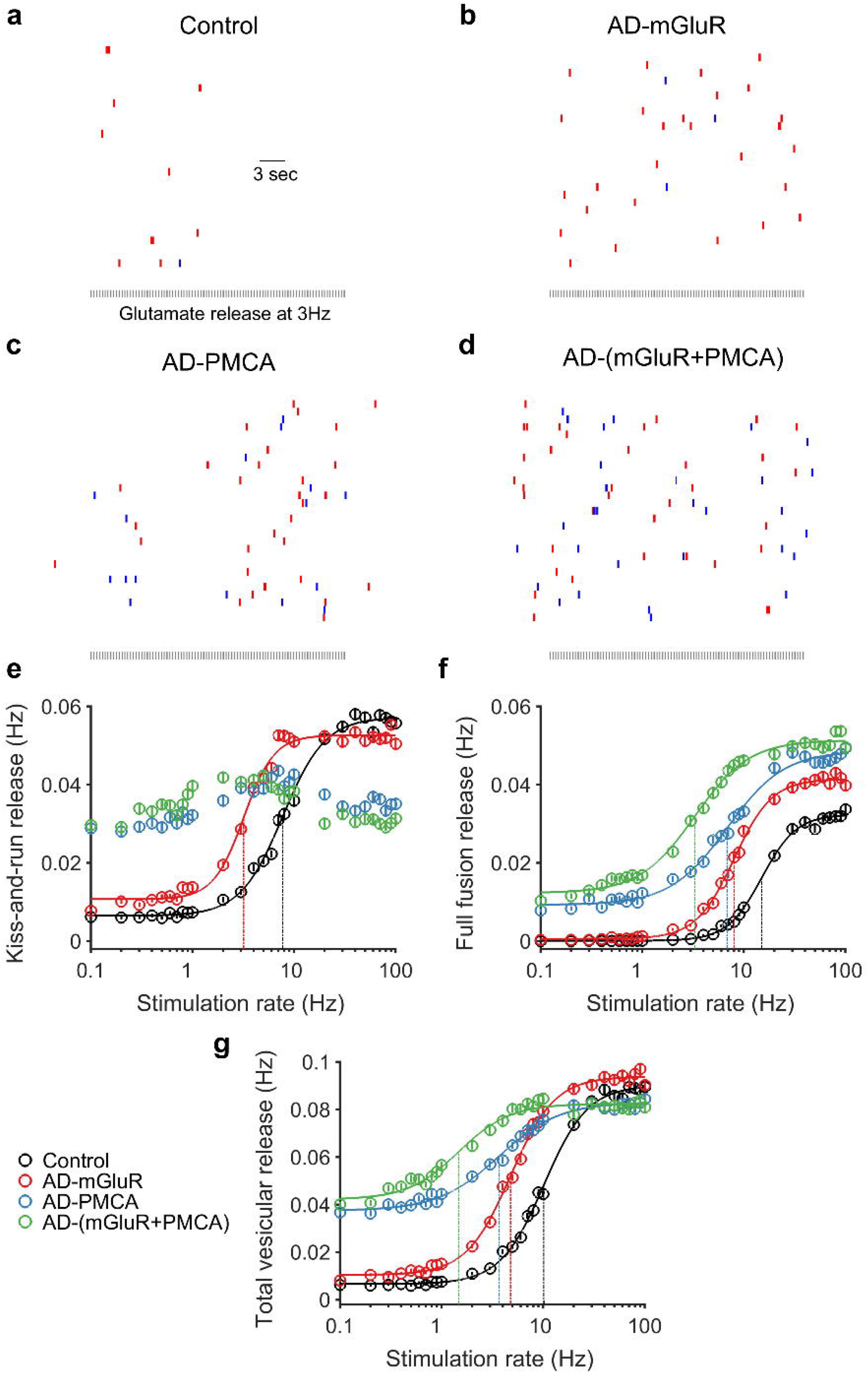
Aβ-induced alteration in PMCA signaling favors full fusion release over kiss-and-run exocytosis. **a-d** Kiss-and-run (red) and full fusion (blue) release events recorded from control and AD astrocytic processes stimulated with spillover glutamate from neuronal release at 3 Hz. **e** Aβ-induced alterations in PMCA signaling abolished the logistic relationship between kiss-and-run release and glutamate stimulation rate. **f** Reduced PMCA activity in AD processes lead to a leftward shift in full fusion release rate. **g** The half maximum of total release rate is lowered in all astrocytic processes with AD pathology. Dotted lines indicate the frequency of half-maximal response for each group.

Both kiss-and-run and full fusion exocytosis were upregulated in the AD-mGluR group compared to control. Accordingly, EC_50_ values obtained by fitting a Hill equation (Eq. 5) to kiss-and-run and full fusion rates were low in the AD-mGluR group compared to control (Figs. 5e & f). The drastic reduction in kiss-and-run exocytosis was very evident in the AD-mGluR group whose EC_50_ value was reduced to less than a third of the control group (Fig. 5e). In contrast, in AD astrocyte process groups with impaired PMCA (AD-PMCA & AD-mGluR+PMCA), the rate of kiss-and-run exocytosis became unresponsive to changes in presynaptic glutamate release rate (Fig. 5e). Both the AD-PMCA groups also had high spontaneous release rates that were mostly kiss-and-run. In the AD-mGluR group, kiss-and-run release dominated at all stimulation frequencies. Whereas in AD groups with inhibited PMCA action, a majority of releases at high stimulation rates (> 10 Hz) were full fusions. AD related abnormalities shifted the relationship between total release rate and stimulation rate leftward with low EC_50_ values in all the AD groups (In Hz, control: 10.2, AD-mGluR: 4.8, AD-PMCA: 3.7, AD-mGluR+PMCA: 1.5) (Fig. 5g).

### Synchrony of Ca^2+^ and release events are enhanced in AD astrocytic processes with altered mGluR pathology

Increased astrocytic synchronization in the form of coherent Ca^2+^ and gliotransmitter release events boosts neural synchrony *in vivo* and *ex vivo*^79,80^. Synchronous Ca^2+^ activity in astrocytes is also associated with pathological neuronal discharges in several disease models including epilepsy and AD^14,81^. We therefore tested synchrony of stochastic Ca^2+^ and release events from AD astrocytic process stimulated with glutamate pulses resembling neuronal release at different rates (0.1-100 Hz). Event synchrony between independent stimulation trials (1000) was computed using the Pinsky-Rinzel algorithm (See Materials and Methods for details).

A polar representation of phase coherence histogram computed for 0.2 Hz stimulation indicates high coherence of Ca^2+^ events in the AD group with enhanced mGluR signaling (Fig. 6a). However, when stimulated with different rates of presynaptic glutamate release, coherence of Ca^2+^ events was high in all AD groups at low rates (< 1 Hz). In contrast, synchrony remained low at all frequencies in the control group (Fig. 6b). These findings on synchronous Ca^2+^ events at a single astrocytic processes is in agreement with previous whole-cell measurements in AD astrocytes *in vivo* (kuchibhotla2008). The model thus revels mGluR signaling, and not PMCA activity, as a cellular substrate for synchronous astrocytic hyperactivity.

**Figure 6.**
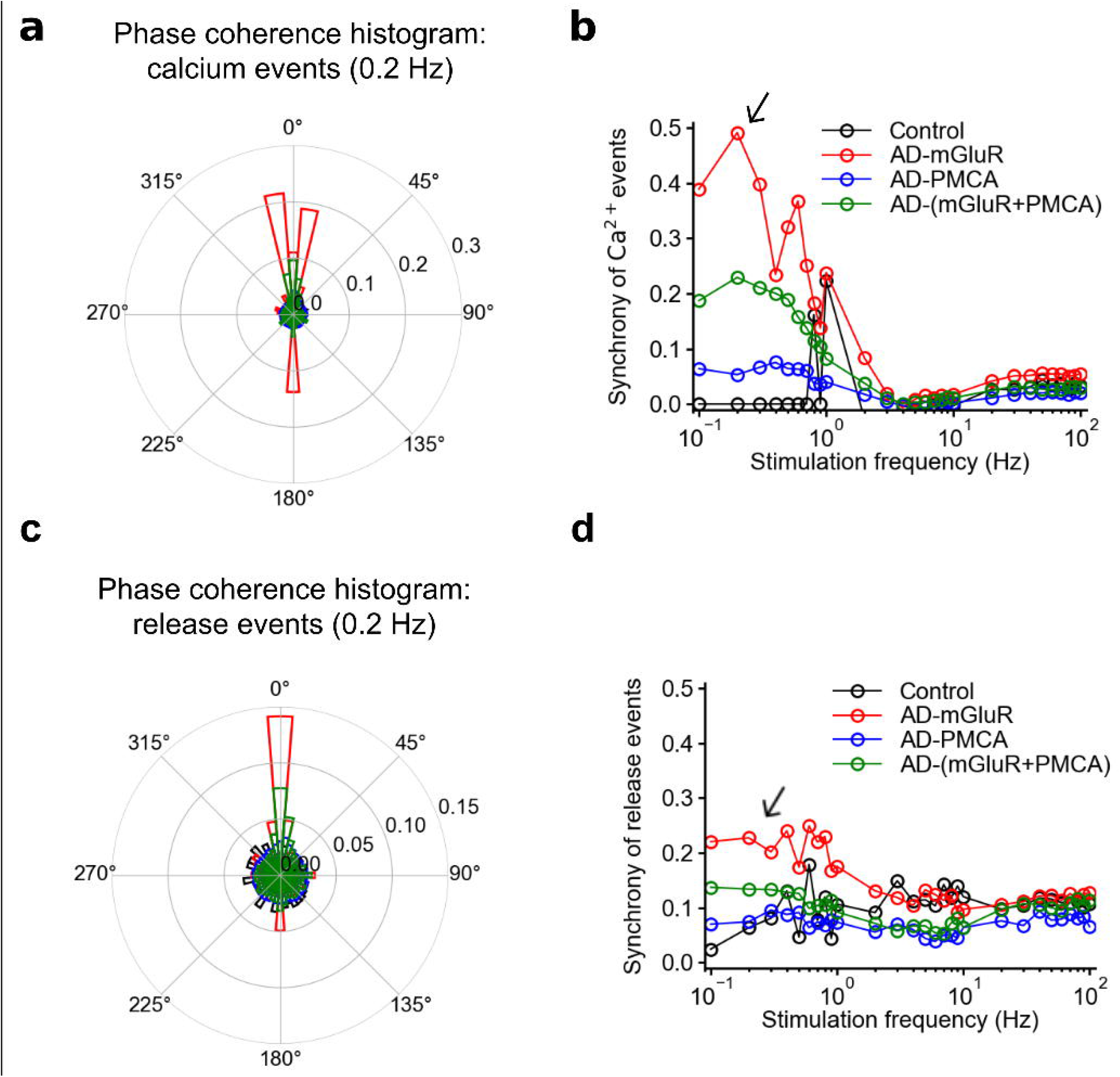
Synchrony (phase coherence) of Ca^2+^events, but not release events, is enhanced in astrocytic processes with AD-related molecular alterations. **a** Polar histogram of phases of Ca^2+^ event from 1000 independent simulation trials of astrocytic processes from control and AD groups that are stimulated with neuronal glutamate release at 0.2 Hz. **b** Ca^2+^ synchrony is high in the AD-mGluR when stimulated at low frequencies. The arrow indicates synchrony for 0.2 Hz stimulation. **c** Polar phase histogram of gliotransmitter release events from 1000 independent astrocytic processes from control and AD groups stimulated with neuronal glutamate release at 0.2 Hz. **d** Synchrony of release events are not much different for the AD groups when compared to control.

Simultaneous measurements of Ca^2+^ and gliotransmitter release events from single astrocytic processes in response to physiological neuronal release is difficult and none exist for AD disease model. Using our systematically validated model we next describe synchrony of gliotransmitter releases in response to astrocytic Ca^2+^ activity (Fig. 6c and d). Similar to what we observed for Ca^2+^ events, coherence of gliotransmitter releases was high in the AD-mGluR group at 0.2 Hz (Fig. 6c). Although, gliotransmitter releases were much less synchronous than Ca^2+^ events in the same trials, at low stimulation rates, synchrony of AD-mGluR group dominated over all the other groups (Fig. 6d).

### Low probability of docked vesicles leads to loss in the temporal correlation between Ca^2+^ and release events in AD astrocytic processes

Apart from synchronizing neuronal ensembles, astrocyte processes also exert synapse-specific control of synaptic transmission^45^. An important prerequisite for this functionality is precise temporal association between Ca^2+^ event arising out of presynaptic release and gliotransmitter release events at a specific tripartite synapse. The close association between calcium activity is clearly seen in experimental data and is reproduced accurately in our model (Figs. 1f and 2f). Following up on our previous results on increased calcium signaling and gliotransmission by AD mechanisms, we next investigated its impact on the temporal coincidence between Ca^2+^ and release events.

Similar to the experimental protocol described previously, we stimulated control and AD astrocytic process with the temporal profile of spillover glutamate at different release rates. Using joint crosscorrelation analysis (JCC), the temporal association between Ca^2+^ and gliotransmitter release events was computed for each stimulation regime (See Model and Methods for details). We first present data from correlation analysis and the nJPSTH matrix computed from 1000 trials of 10 Hz stimulation train for control and AD groups (Fig. 7a-d).

**Figure 7.**
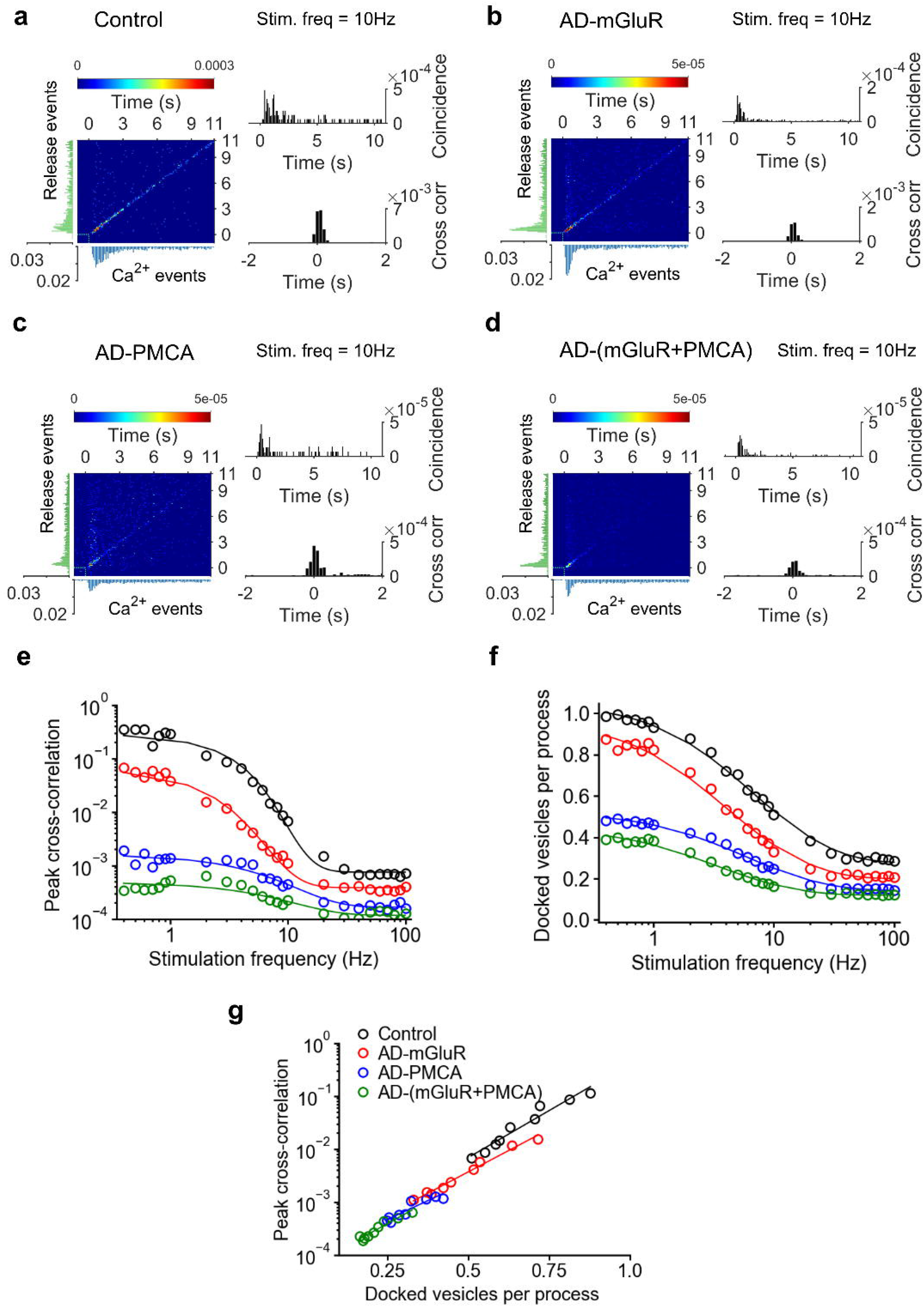
Temporal relationship between Ca^2+^ and release events is reduced in astrocytic processes with AD pathology. **a** nJPSTH (*left*), coincidence (top) and cross correlation (*bottom*) histograms between Ca^2+^ and release events computed from 1000 independent simulation trials of the control astrocyte process stimulated with neuronal glutamate release at 10 Hz. At the bottom and left side of the nJPSTH are PSTHs for Ca^2+^ and gliotransmitter release events. **b-d** nJPSTH, coincidence and cross correlation time histograms of AD-mGluR, AD-PMCA and AD-(mGluR+PMCA) groups. **e** Peak cross correlations between Ca^2+^ and release events at different stimulation rates are low in all AD astrocytic processes. **f** The average number of docked vesicles computed from 1000 independent trials is low in all AD groups. **g** The linear relationship between peak cross correlation and docked vesicles at the theta stimulation range (2-10 Hz) indicates rapid vesicle depletion in AD processes as a mechanism for the loss in temporal association between Ca^2+^ and release events. Fits to Hill equations are in solid lines.

As evident from the colored representation of the nJPSTH matrix, in the control group, Ca^2+^ and gliotransmitter release events were highly coincident for more than 10 s from the stimulus onset (Fig. 7a, *left*). This was further confirmed from the coincidence histogram computed from the diagonal elements of the nJPSTH matrix (Fig. 7a, *top-right*). In agreement with experimental findings^22^, cross-correlations obtained from para-diagonal bins of nJPSTH matrix peaked at 100 ms lag (Fig. 7a, *bottom-right*).

However, compared to the control group, nJPSTH matrices of AD groups indicated lower coincidence between Ca^2+^ and gliotransmitter release events (Fig. 7a-d, *top-left*). Evidently, in all the AD groups, coincidence was low and decayed sharply after the stimulus onset (Fig. 7a-d, *top-right*). All the AD groups also had low and broadened cross-correlations values (Fig. 7a-d, *bottom-right*).

We next extended the cross-correlation analysis at a single astrocytic process to low and high stimulation frequencies (0.4-100 Hz). Similar to what we found with Ca^2+^ synchrony, peak cross-correlation was high in all the groups when the frequency stimulation was low. However, in stark contrast, Ca^2+^ and vesicular release events were less correlated in all the AD groups. Fits to a logistic function (Eq. 9) also yielded lower values in all the AD groups compared to the control group (Fig. 7e).

To further understand the reason behind this surprising result, we quantified average number of vesicles docked during stimulus presentation. Clearly, docked vesicles decreased with higher stimulation rates and was low in all the AD groups (Fig. 7f). Since docked vesicles mediate fast and synchronous kiss-and-run exocytosis, we next examined its link with peak cross-correlation between Ca^2+^ and gliotransmitter release events when stimulated by glutamate pulses delivered at the theta frequency range (2-10 Hz) (Fig. 7g). The clear linear relationship between the two further imply rapid depletion of docked vesicles as the underlying mechanism for the loss in temporal coincidence between Ca^2+^ and release events in AD astrocytes.

## Discussion

While AD clearly manifests as a synaptic abnormality early on, little is known about the contribution by astrocytes to its pathogenesis^10,82^. To address this gap, we harnessed experimental data across multiple-scales and developed a biophysical model for astrocytic calcium response and ensuing gliotransmitter release at a typical CA3-CA1 synapse. The model is strongly rooted in extant experimental evidence and comprises of a realistic description of molecular components such as membrane receptors, local ER extent, vesicle types and importantly, the synaptotagmins. The model accurately captures statistical distributions, timescale and amplitudes of both Ca^2+^ and gliotransmitter release events as reported experimentally^22^. We next characterized both Ca^2+^ signaling and vesicle release at a single process to show that both synchrony and rate of astrocytic Ca^2+^ responses to trains of presynaptic glutamate release are enhanced in AD. The most crucial insight that arises out of this study is the observed loss in temporal precision of individual Ca^2+^ and vesicle release events in all the AD astrocytic groups. The model predicts impaired mGluR and PMCA signaling to distinctly enhance Ca^2+^ synchrony and spontaneous hyperactivity, respectively. Together, our results provide novel mechanistic insights on AD-related astrocytic dysfunction as well as suggest impaired astrocytic feedback at tripartite synapses as a critical functional consequence of AD pathology.

A wealth of whole-cell Ca^2+^ imaging and computational studies have linked IR_3_R-mediated slow calcium waves in astrocytes and long-range synchronization of neuronal networks^34,80,83,84^. The traditional view that astrocytes primarily impart slow modulation has been challenged recently by studies that reported high amplitude Ca^2+^ transients and Ca^2+^-mediated release from astrocytic processes^22,85–87^. Despite the indication that astrocytic processes can rapidly modulate synaptic transmission, there is no clear understanding of the biophysical mechanisms that mediate the tight temporal correspondence between Ca^2+^ events and vesicular release at a single process^22,45,88,89^. From the model it is clear that increased presence of ER, the high Ca^2+^ levels in PAPs, rapid opening of IP_3_Rs, low cytoplasmic buffering, a presence of a fast Ca^2+^ sensor and rapid vesicle recycling together orchestrate the tight temporal feedback at tripartite synapses.

Based on experimental evidence on Ca^2+^ compartmentalization, we treated each astrocytic process as an independent unit with high (50%) ER-to-cytosol volume ratio^90^. Calcium responses at these tiny domains are further constrained to tight subcellular domains due to diffusion constraints that increase the influence of mGluR_5_ and limits the spatial spread of IP_3_ within individual processes^21,91^. The presence of multiple Ca^2+^ regulating mechanisms and a small cluster of IP_3_Rs leads to stochastic Ca^2+^ events in astrocytic processes similar to Ca^2+^ puffs observed in other cell types^70,92^. Our stochastic model with a single cluster of five IP_3_Rs accurately reproduced both temporal characteristics and the distribution of Ca^2+^ event amplitudes, rise times and decay constants as reported by previous studies^22–25^. The model also reproduced several critical determinants of mGluR-mediated Ca^2+^ events, such as mGluRs dose-response, ER Ca^2+^ levels (400 mM) and SERCA refilling time (100 s)^49,93–96^. Additionally, single process Ca^2+^ responsiveness to glutamate pulses from the model is also in good agreement with somatic Ca^2+^ imaging data recorded from astrocytic cultures^4^. Lastly, we compared model results to the temporal histogram of Ca^2+^ events from entire astrocytes^22^. Based on our estimate on the number of processes within the imaging volume of a single astrocyte, we pooled data from 400 independent simulation trials. While it is likely that there is variability in intrinsic parameters (IP_3_ receptors, geometry etc) across astrocytic processes, our simulations account for more than 90% of all the events observed experimentally. These independent validations across a wide range of experimental data indicate that the biophysical model developed here is an accurate representation of Ca^2+^ signaling in astrocytic processes.

An important functional consequence of astrocytic Ca^2+^ events is gliotransmitter release. Indeed, both kiss-and-run and full fusions exocytosis with distinct timescales of exocytosis, endocytosis and reacidification have been reported in astrocytic processes^22,27^. This evidence supports two separate molecular pathways for vesicular release and recycling. Alongside two synaptotagmins (*Syt4* & *Syt7*) that differ in reaction kinetics and Ca^2+^ sensitivity, astrocytic processes are also thought to harbor docked and mobile vesicles^22,28,29,55,56,97^. Activation of *Syt4* by fast and high amplitude Ca^2+^ events near the ER tubuli which is tightly juxtaposed to the plasma membrane near docked vesicles is thought to result in rapid kiss-and-run release^22^. On the other hand, *Syt7* whose Ca^2+^ affinity is high can be triggered by low amplitude and slow Ca^2+^ events to result in slow asynchronous release of mobile vesicles. Indeed, EM studies also report that at astrocytic processes, vesicles could be seen as far as 200 nm away from the plasma membrane and a fraction of them are suggested to be mobile^5,56^. Considering the very limited spatial extent of a single process where only a few vesicles are docked, the presence of mobile vesicles provides an alternate pathway for experience dependent modulation of release kinetics. Notably, the proportion of slow full-fusion and fast transient releases in astrocytes is stimulus dependent^27,32^. In agreement to the above, we observed a shift from kiss-and-run to full fusion exocytosis in the AD astrocytic process.

In order to model AD astrocytes, we considered two well known molecular signatures of Aβ accumulation that are critical for astrocytic Ca^2+^ regulation, namely, dysfunctional PMCAs and elevated mGluR signaling^13,70–73^. Our study provides several clear and experimentally testable predictions that relate Aβ-induced alterations with astrocytic Ca^2+^ signaling at a single process level. Firstly, our prediction that AD astrocytic process have spontaneous Ca^2+^ activity is consistent with experimental reports from AD animals and *ex vivo* A β treatment^11,14,78,98^. Our study specifically associates reduced PMCA activity with increased cytosolic resting Ca^2+^ and enhanced spontaneous Ca^2+^ activity. Again, increased resting intracellular Ca^2+^ has been reported *in vivo* in AD astrocytes, although the mechanism was not known^14^. Secondly, results from the model indicate enhanced mGluR signaling as the underlying molecular mechanism for Ca^2+^ event synchrony and similar to this finding, synchronous bulk Ca^2+^ signals are found in AD astrocytes^14^.

Despite the well-established evidence for abnormal Ca^2+^ signaling in astrocytes, there is no clear understanding of the consequences of abnormal calcium signaling on gliotransmitter release at a single process. We show that kiss-and-run and full fusion exocytosis mediated by *Syt4* and *Syt7,* respectively, were differently affected by mGluR and PMCA mechanisms. Enhanced mGluR signaling increased both kiss-and-run and full fusion releases as well as their inter-trail synchrony when stimulated with trains of glutamate pulses mimicking presynaptic activity. In contrast, PMCA alteration abolished both synchrony and the logistic relationship of kiss-and-run release with stimulation strength.

Lastly, our study uncovered a very unique dependence between AD-related enhanced Ca^2+^ dynamics and loss in temporal precision of gliotransmitter release at a single process. Unlike in control astrocytes whose release events were concurrent with Ca^2+^ events within a time window of 100 ms, all the AD groups displayed a loss in this precision very early on after the stimulus onset. We also report a clear linear relationship between this loss in temporal coincidence and rapid depletion of docked vesicles when stimulated in the theta frequency range (2-10 Hz) which also corresponds to the dynamic range of astrocytic responses at a single process. Thus when affected by AD pathology, astrocytic processes become more synchronous but lose their ability to provide timely feedback at single tripartite synapses^99^. As suggested by previous computational and experimental studies, both these mechanisms can affect how astrocytes modulate presynaptic release probability and synaptic information transfer in the hippocampus^1,89^.

In summary, we incorporated diverse experimental data to construct a biophysically detailed model for calcium signaling and the corresponding gliotransmitter release at a single astrocytic process. The model accurately reproduces both the kinetics and statistical profiles of both calcium and release events as reported by previous experiments. The biophysical model we describe is firmly grounded in physiology and is flexible to simulate diverse experimental setting to predict novel astrocytic mechanisms on synaptic signaling. The difficulty in performing actual experiments at the highly ramified astrocytic processes further enhance the value of the model. Based on our model we show that in AD astrocytes, enhanced depletion of docked vesicles leads loss in correspondence between calcium events and gliotransmitter release events. We hypothesize that this mismatch in information transfer at a crucial hippocampal tripartite synapse may contribute to higher level cognitive deficits associated with AD.

## Data availability

All the codes of our model and simulations will be made publicly available at GitHub repository (https://github.com/anupgp/astrocytic_process) after publication.

## Acknowledgements

AGP was supported by a Postdoctoral Fellowship from Indian Institute of Science Education and Research Pune (IISER-P/Ext/PDRF/AP-20145047/10/2016) and Wellcome Trust/DBT India. S.N. was funded by Wellcome Trust/DBT India Alliance (IA/I/12/1/500529) and Indian Institute of Science Education and Research Pune.

## Author Contributions

A.G.P. and S.N. designed the research and wrote the paper. A.G.P. wrote the codes to build the model, perform the simulations and analyzed the data.

