## Supplemental Figures for "Amyloid pathology disrupts gliotransmitter release in astrocytes"

Supplementary Materials

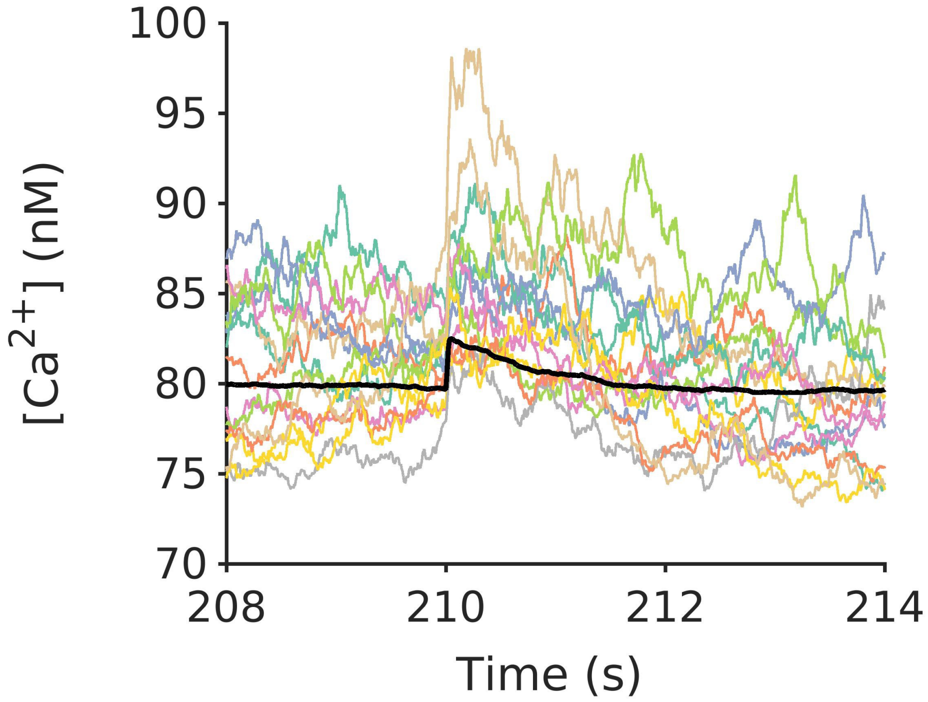


**Figure 1.** Overlay of calcium events from an astrocytic process stimulated with the glutamate profile from a single release event. Each color represents $Ca^{2+}$ response from a single trial that responded to single vesicle glutamate stimulation. Due to the stochastic nature of $Ca^{2+}$ responses, not every trial leads to a clearly identifiable response. The black trace represents the average from roughly 20 responses.

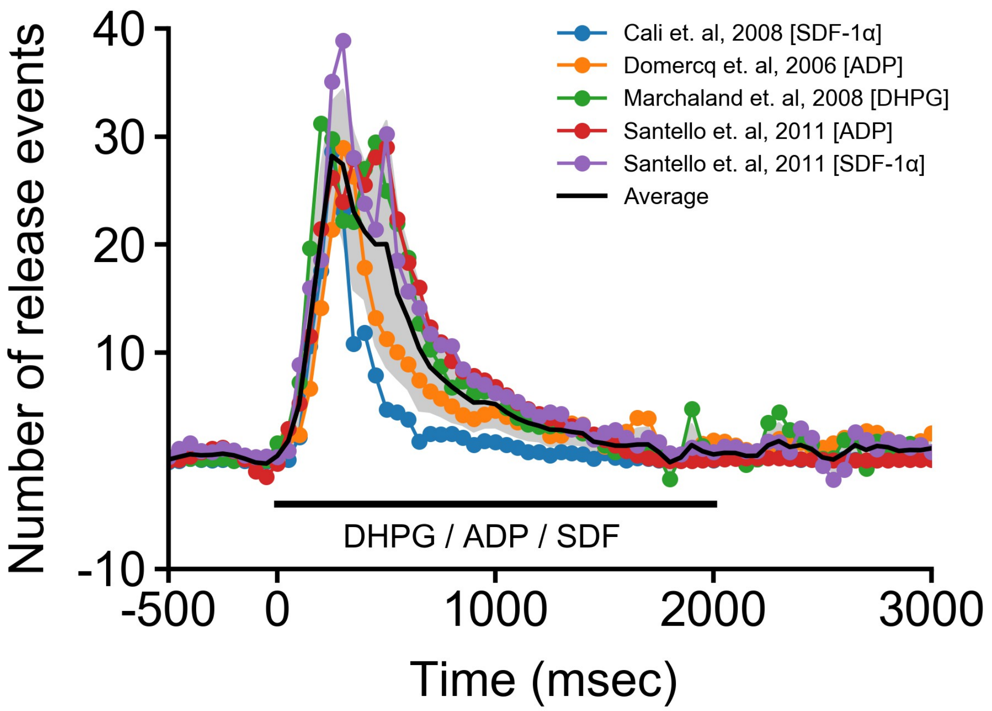


**Figure 2**. Results reproduced from published studies that have monitored vesicular release from single astrocytes. The studies have used Total Internal Reflection Microscopy (TIRF) to count vesicles released in the evanescent field (~ 90 nm) from entire astrocytes expressing pH sensitive fluorescent dye on glutamate containing vesicles (VGLUT1-pHluorin). Astrocytes were stimulated with saturating concentrations of various agonists (2 sec) to release $Ca^{2+}$ from internal stores that lead to subsequent $Ca^{2+}$-dependent vesicular release.

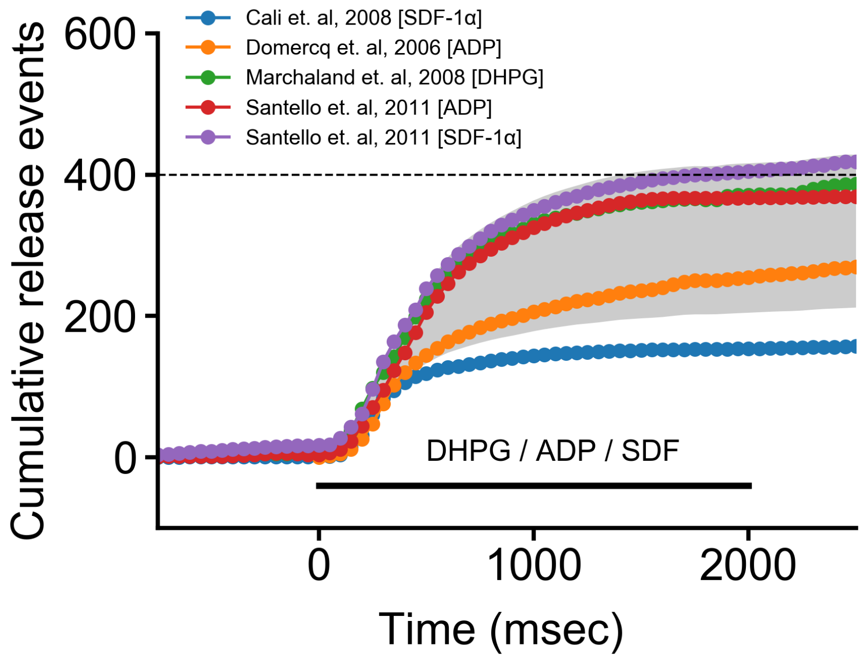


**Figure 3**. Results reproduced from published studies on the cumulative count of released vesicles from astrocytes. Astrocytes expressed pH sensitive fluorescent dye on glutamate containing vesicles (VGLUT1-pHluorin) and were stimulated with saturating concentrations of various agonists (2 sec) to release $Ca^{2+}$ from internal stores that lead to subsequent $Ca^{2+}$-dependent vesicular release.

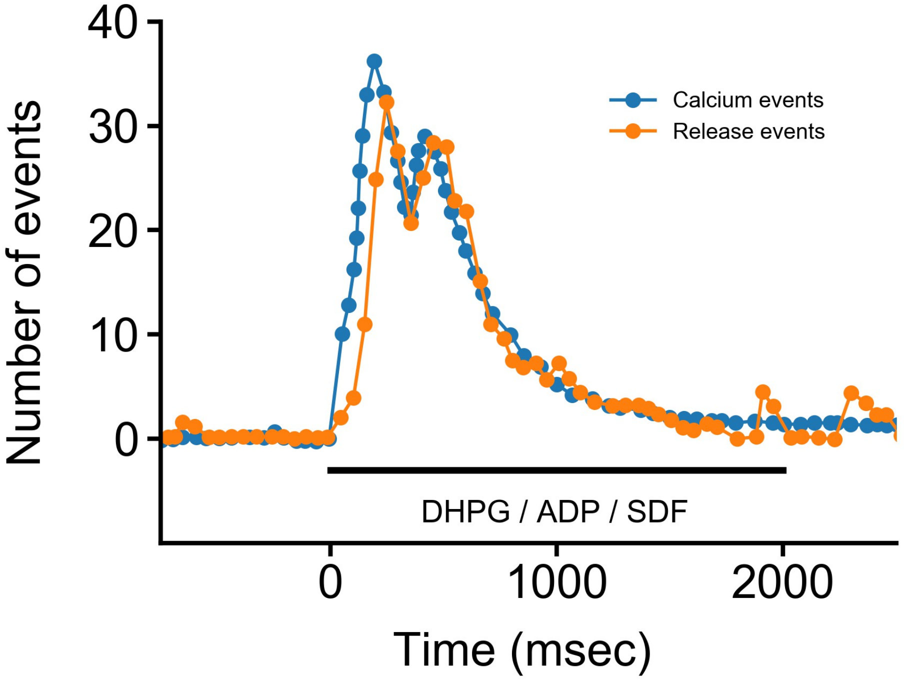


**Figure 4**. Results reproduced from Marchaland et al., 2008 on the number of $Ca^{2+}$ and release events from microdomains of entire astrocytes. Astrocytes expressing VGLUT1-pHluorin were imaged using TIRF microscopy. A tight match was evident between $Ca^{2+}$ and release events on their temporal profiles following mGluR1 stimulation with a saturating concentration of DHPG (2 sec, 100 μM).

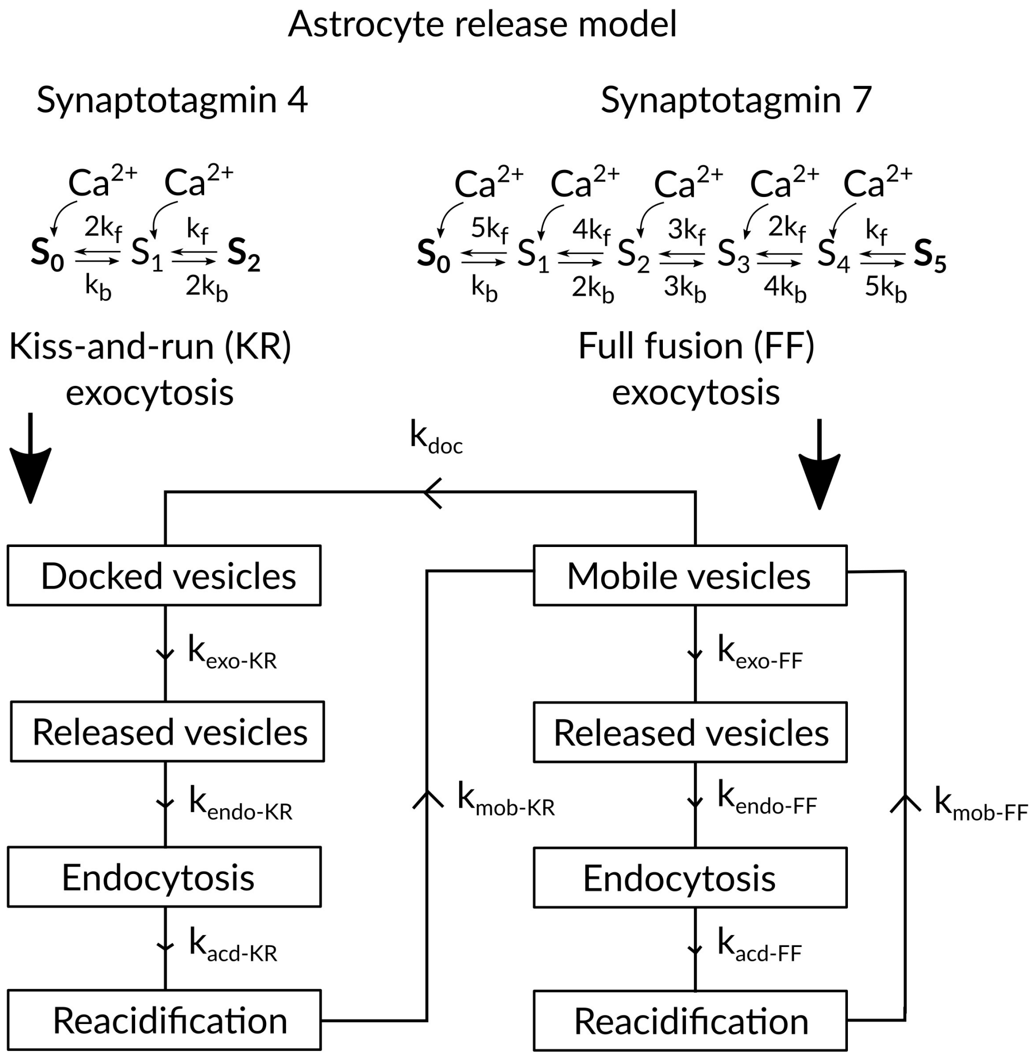


**Figure 5**. A detailed schematic of the gliotransmitter release model. Each astrocytic process contained two $Ca^{2+}$ sensors (synpatotagmin 4 & 7) that independently mediated kiss-and-run and full fusion exocytosis from two seperate vesicle pools. Following kiss-and-run release, docked vesicles are endocytosed and reacidifed before being taken up into the mobile vesicle pool. Full fusions arise from the activation of *Syt7* and exclusively from the mobile vesicle pool. Docked vesicles are finally replenished in a rate dependent manner from the mobile vesicle pool.
